# Subsidy Accessibility Drives Asymmetric Food Web Responses

**DOI:** 10.1101/2020.11.09.374629

**Authors:** Marie Gutgesell, Bailey C. McMeans, Matthew M. Guzzo, Valesca deGroot, Aaron T. Fisk, Timothy B. Johnson, Kevin S. McCann

## Abstract

Global change is fundamentally altering flows of natural and anthropogenic subsidies across space and time. After a pointed call for research on subsidies in the 1990s, an industry of empirical work has documented the ubiquitous role subsidies play in ecosystem structure, stability and function. Here, we argue that physical constraints (e.g., water temperature) and species traits can govern a species’ accessibility to resource subsidies, and that these physical constraints have been largely overlooked in the subsidy literature. We examined the input of a high quality, point-source anthropogenic subsidy into a recipient freshwater lake food web (i.e., released net-pen aquaculture feed in Parry Sound, Lake Huron), to demonstrate the importance of subsidy accessibility in governing recipient whole food web responses. By using a combined bio-tracer approach, we detect a gradient in accessibility of the anthropogenic subsidy within the surrounding food web driven by the thermal tolerances of three constituent species. This thermally-driven accessibility gradient drives asymmetrical changes in food web structure, effectively rewiring the recipient lake food web and altering patterns in secondary production with yet unknown stability consequences. Since aquaculture is predicted to increase significantly in coming decades to support growing human populations, and global change is altering temperature regimes, then this form of food web alteration may be expected to occur frequently. We argue that subsidy accessibility is a key characteristic of recipient food web interactions that must be considered when trying to understand the impacts of subsidies on ecosystem stability and function under continued global change.

## Introduction

Ecosystems are intrinsically connected through space and time by flows of energy (i.e., subsidies) that ultimately help govern the receiving ecosystem’s stability and function. Foundational work by empirical ecologists in the late 1990’s demonstrated the ubiquity of natural subsidies (e.g., nutrients, detritus, prey) connecting landscapes across a broad range of spatial and temporal scales (Polis & Strong, 1996; Polis & Winemillar 1996; Polis et al., 1997; Sears et al., 2004). This empirical work led to theory demonstrating that subsidies are pivotal in determining the stability and community composition of recipient food webs (Polis et al., 1997; Huxel & McCann, 1998; Sears et al., 2004, Takimoto et al. 2002). While subsidies occur naturally, a growing human population is increasing the prevalence of anthropogenic subsidies to natural ecosystems (e.g., agricultural nutrient run-off, sewage, net-pen aquaculture waste). Akin to natural subsidies, anthropogenic subsidies also have the potential to influence recipient food web dynamics (DeBruyn et al., 2004; Rodewald et al., 2011; Newsom et al., 2015; Singer et al., 2016; Lee et al., 2018; Johnson et al., 2018; DeBruyn et al., 2020). As food web dynamics govern whole ecosystem stability and function (de Ruiter et al., 1995; Neutel et al., 2002), it is imperative to understand how recipient food webs respond to both natural and anthropogenic subsidies under continued global change.

It is now well recognized that fluxes of subsidies play a major role across most ecosystems (Leroux & Loreau, 2008) and also ought to significantly impact the stability and functioning of recipient ecosystems (Huxel & McCann, 1998; Takimoto et al., 2002; Leroux & Loreau, 2008). An industry of empirical and theoretical research has emerged since Polis and others introduced subsidies to food web theory in the 1990s with researchers exploring the various ways subsidies can impact recipient ecosystems. Research has concentrated on: i) subsidy quality and quantity; ii) the temporal nature of subsidies (e.g., Nakano et al., 1999; Nakano & Murakami, 2001; Sears et al., 2004); iii) the role of subsidies in driving trophic cascades (Huxel & McCann, 1998), and; iv) the roles space and ecosystem type play in governing subsidy flow rate, assimilation, and magnitude of resultant impacts on food web dynamics (Leroux & Loreau, 2008). While this research has made major progress, surprisingly little work has looked at the role of species traits (e.g., thermal preference) and physical habitat characteristics (e.g., water temperature) in determining subsidy accessibility, and how this may govern whole food web responses. Since accessibility may allow us to better understand how subsidies integrate into whole recipient webs this is a major gap in subsidy research.

While subsidies are generally thought to be ubiquitous in nature, it is less understood how the physical location of a subsidy within a habitat or ecosystem impacts its ability to be accessed and thus assimilated by members of the local food web. Subsidy accessibility, defined as the degree of availability of subsidies to recipient species based on their physiological (e.g., thermal tolerance) and physical habitat limitations, can drive differential increases in productivity throughout the receiving food web. According to optimal foraging theory, if capable, species will alter their foraging behaviour to access the highest density/quality resource available (i.e., also referred to as the Birdfeeder Effect at the community level (Eveleigh et al., 2007)). However, species may be limited in their ability to respond to these changes in local resource densities by traits, such as mobility and thermal tolerance. For example, subsidies entering the nearshore zone of a temperate lake may be expected to be most easily accessed by warm adapted fishes and least accessible to cold adapted species. Therefore, in the example above, we may expect to see the greatest alteration in diet towards subsidies in warm water species and reduced or absent alteration in diet in cold water species. This type of asymmetrical subsidy accessibility and uptake can in turn drive differential changes in productivity and thus food web structure throughout whole recipient ecosystem. Here, we combine the ideas that species optimally forage within their physical and physiological capabilities and the asymmetric impacts of spatial subsidies on receiving food webs, to suggest that species’ ability to access subsidies can drive differential (or asymmetric) changes in food web productivity and structure.

In this study, we broadly examine the dispersal of a point-source anthropogenic subsidy into a temperate lake food web that contains dominant mobile, generalist top predators from each of the three thermal guilds (i.e., a gradient in trait responses from cold-to coo-l to warm-water fish) (Magnuson et al., 1990). We build off empirical evidence of assimilation of a high-quality anthropogenic subsidy in our study system, released net-pen aquaculture feed (Johnson et al., 2018), to demonstrate how accessibility to this subsidy is key in governing the receiving food web’s response to this novel anthropogenic energy source. Differential accessibility throughout the recipient food web has the potential to differentially alter productivity and thus asymmetrically alter food web structure (e.g., differential changes in food chain length). As changes in food web structure can drive alterations in food web dynamics, this is a key first step in understanding how subsidy accessibility may ultimately influence food web stability. Here, we use a combined bio-tracer approach to trace the fate of the anthropogenic subsidy throughout the lake food web, and then combine hydroacoustic, fish biomass, and stable isotope data to detect asymmetrical shifts in food web productivity and structure (i.e., food chain length) across a thermal gradient.

## Methods

### Site Description

Parry Sound, located in Georgian Bay of Lake Huron, was chosen as our study site to investigate how accessibility to a point-source anthropogenic subsidy influenced assimilation into a surrounding food web and alterations in subsequent food web productivity and structure. This location presents an ideal study system to address this question as there is a large consistent, long-term, high quality point-source anthropogenic subsidy (i.e., released net-pen feed & waste from Aqua-Cage Fisheries Ltd.). Rainbow trout are reared within the Aqua-Cage net-pens and are fed daily from feed boats that move between cages from ~9:00-16:00 each day (Kana Upton, Aqua-Cage Fisheries Ltd., Pers. Comm). Feed consists of pelleted food manufactured from both aquatic and terrestrial energy sources (see Appendix1: Table S1 and Table S2 for composition of the fish feeds). Parry Sound undergoes thermal stratification during summer, which creates a natural thermal gradient from the near-shore, warm littoral zones (that are more accessible to fishes with warmer thermal preferences) to the off-shore, cold pelagic zones (that are more accessible to fishes with colder thermal preferences). The location of the point-source subsidy in the deep, cold region of the lake thus sets up a natural gradient of accessibility to the net-pen feed from most accessible (cold-water species), intermediate accessibility (cool-water species), to least accessible (warm-water species).

To determine if any changes in diet and food web structure in Parry Sound were in fact driven by differential assimilation of net-pen feed, we selected multiple control sites around Georgian Bay, Lake Huron (see Appendix 1: Figure S1 for map). Control sites were selected by the following criteria: no presence of off-shore point-source anthropogenic subsidy, far distance to ensure no mixing of populations between subsidy and control sites, and presence of comparable species. One control site was sampled in 2016 (Shawaniga/Sturgeon Bay) and three control sites were sampled in in 2017 (Shawniga/Sturgeon Bay, Key Harbour, and Dyers Bay).

### Fish & Baseline Sampling

Muscle tissue samples were collected from fish and whole invertebrates for stable isotope (SI) and fatty acid (FA) analysis from both Parry Sound and control sites in 2016 and 2017. Fish and invertebrate species were selected to recreate a tri-trophic generalist lake food web that contains a dominant generalist top predator in all three thermal guilds (lake trout (cold), walleye (cool), smallmouth bass (warm)), cold off-shore (alewife & rainbow smelt) and cool near-shore (yellow perch) forage fish, and off-shore/near-shore baseline invertebrates. Figure 1 shows conceptual diagram of the tri-trophic generalist lake food web sampled.

**Figure 1.**
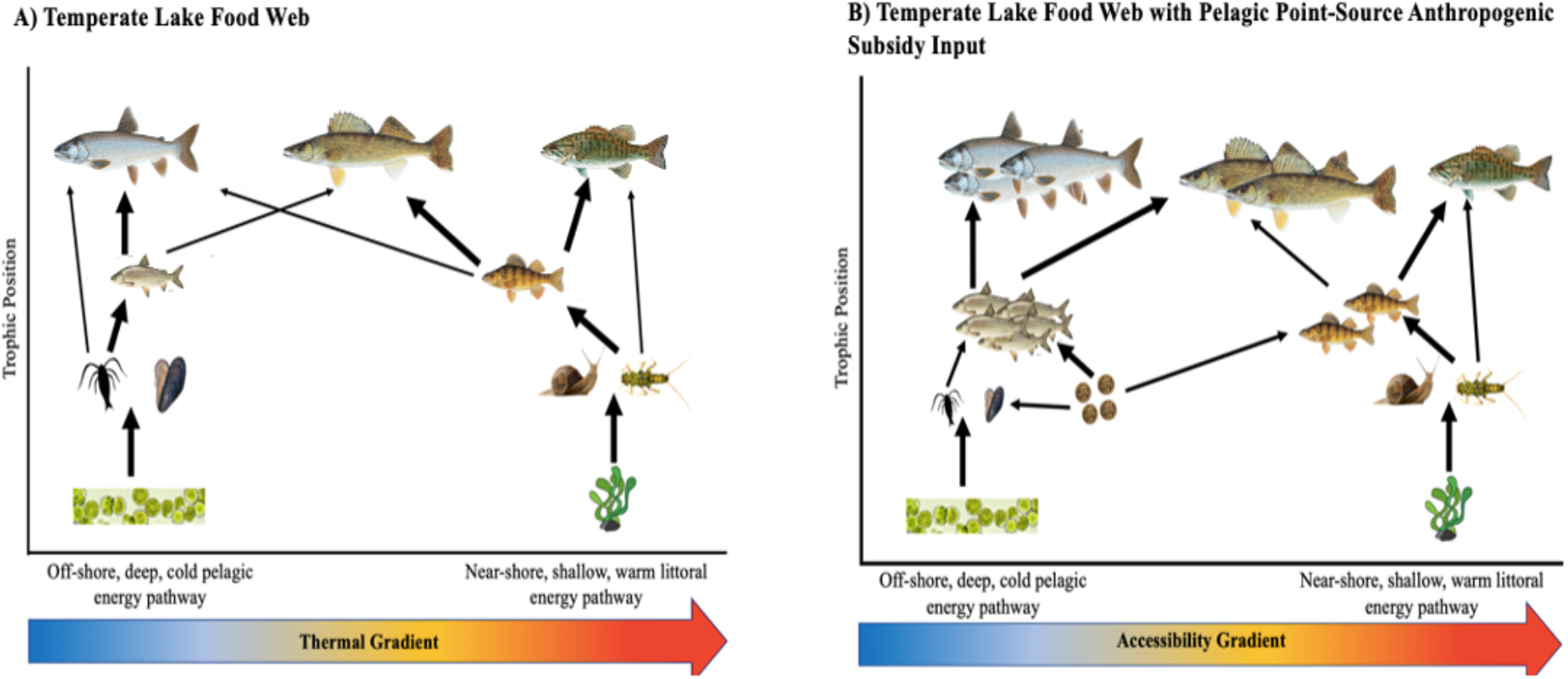
Conceptual figure of food web interactions within (a) natural temperate lake food web and (b) a temperate lake food web with a pelagic point-source anthropogenic subsidy input, which here is released net-pen feed. It is predicted that net-pen feed will alter food web structure through asymmetric accessibility to surrounding food web. Therefore, we expect to see strong assimilation of net-pen feed in cold water fish and subsequent increases in trophic level and biomass. It is expected that some assimilation occurs in cool water fish, and no assimilation of net-pen feed occurs in warm water guild.

Fish samples were collected using Ontario multi-mesh gill nets following a modified broad-scale monitoring (BsM) protocol of the fish community (Sandstrom et al., 2013) and supplemented with targeted angling. Both overnight and daytime sets were employed in the sampling protocol. All gill nets were set for a duration of ~12 h before retrieving fish. Weight (g), length (mm), and two muscle tissue plugs from behind the dorsal fin were collected from fish. Non-predatory insect larvae (i.e., *Ephemeroptera* larvae), snails and mussels (*Unionidae sp.,* and *Dreissena polymorpha*) were collected from each sampling site to provide baseline stable isotope values. Additionally, feed and muscle tissue from farmed rainbow trout was also collected each year Aqua-Cage Fisheries in Parry Sound. All samples were stored at −20°C after collection until further analysis.

One muscle tissue plug from all fish, feed, and baseline samples collected were prepared for stable isotope analysis. Samples were thawed in the lab and dried at 60 °C in a drying oven for 48 h. Once dried, they were individually ground using a mortar and pestle and scooped into a labelled centrifuge tube. The samples were sent to the University of Windsor GLIER (Windsor, ON, Canada) laboratories for carbon and nitrogen isotopic analysis. Muscle tissues from a subset of the fish collected were sent for fatty acid analysis, along with feed and baseline samples. Frozen samples were delivered to Ryerson University – The Arts Lab (Toronto, ON, Canada) in 2016, and both the Laboratory of Aquatic Sciences (Chicoutimi, QC, Canada) and Lipid Analytical Services (Guelph, ON, Canada) in 2017. All labs conducted fatty acid analysis using a combination of Bligh & Dyer and Morrison & Smith methods (Bligh & Dyer, 1959; Morrison & Smith, 1964). Individual FA weights (ug/g) were converted to a % FA composition and fatty acids with >1% presence were retained for analysis.

### Detecting Assimilation of Net-Pen Subsidy into Surrounding Food Web

To determine which fatty acids were key indicators of net-pen aquaculture feed (i.e., fatty acids that are known to be primarily dietary-derived and are significantly higher or lower in the feed than in natural surrounding prey sources) a principal components analysis (PCA) was conducted on fatty acids with >1% average proportions present in forage fish, feed, and baseline fatty acid profiles from all sites. Fatty acid % were standardized to a mean of zero and unit variance prior to their inclusion in the PCA, and separate PCAs were performed for 2016 and 2017 because different fish feeds with different compositions were used in each year (see Supplementary Table 1, Table 2, & Figure 2). Fatty acids with significant loadings (> +/−0.3) along PC1 (indicating significantly higher amounts in feed than natural food sources) were retained as Feed Indicator Fatty Acids (FIFA).

**Figure 2.**
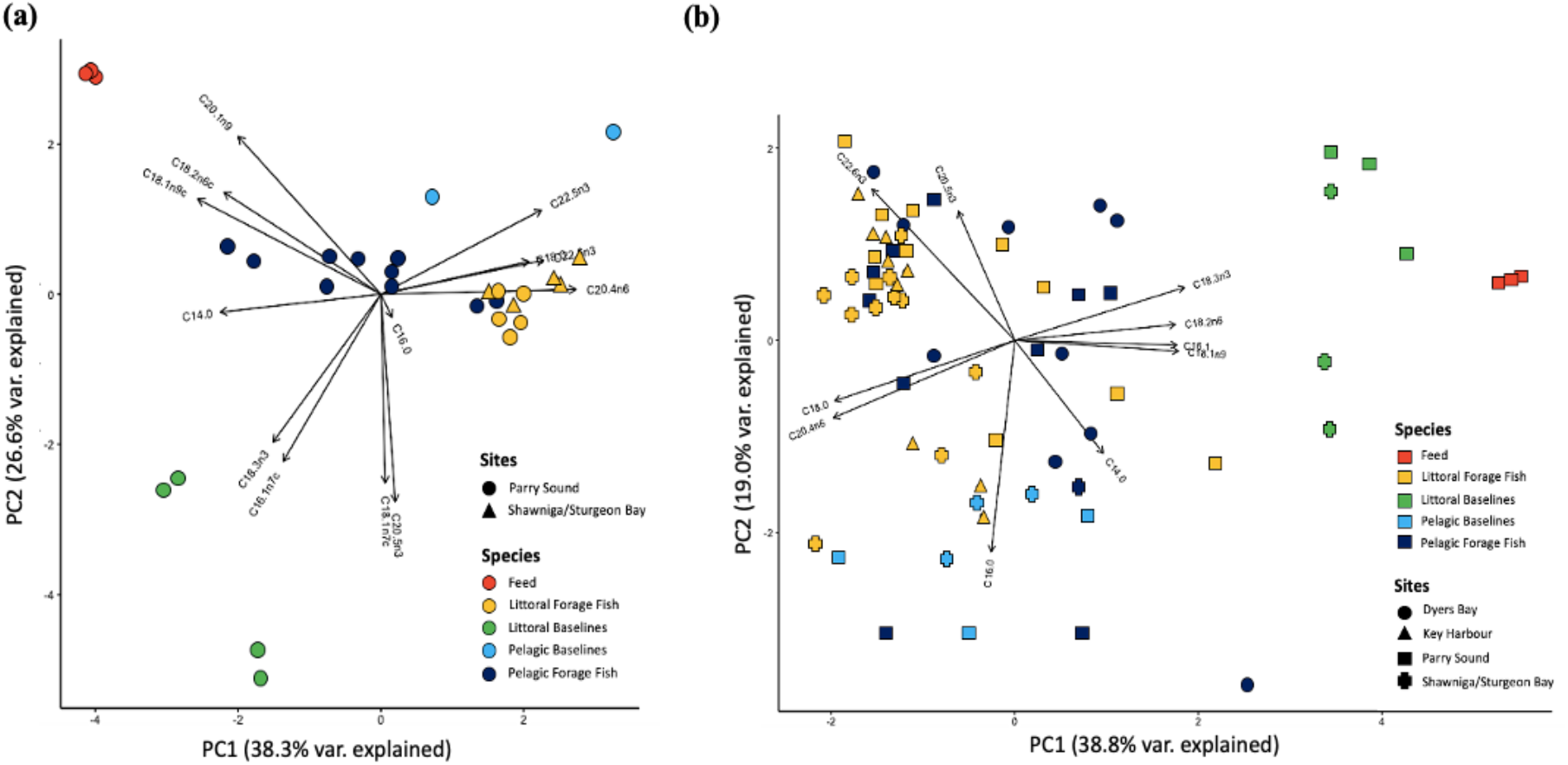
Principal component analysis of fatty acids >1% of total fatty acid composition of all prey fish, baseline, and feed samples analyzed in (a) 2016 and (b) 2017. Feed separated from lake biota (except for littoral baselines) along PC1 in both 2016 and 2017. 2016 feed was separated by 3 FA (higher levels of 18.2n6, 18.1n9, 20.1n9). In 2017, feed was separated by 4 FA (higher levels of 18.2n6, 18:3n3, 18.1n9, and 16:1n7).

FIFA composition of each individual species was then compared between net-pen and control sites using PCAs. The goal of this analysis was to determine if there was distinct clustering of net-pen and control populations driven by the FIFA. To test whether the distance between the net pen and control sites increased from warm to cool to cold water species (reflecting increasing access to and assimilation of the net-pen subsidies), we compared the Euclidean distance between the centroids of the clusters within the PCA. PERMANOVA was used to test for significant differences in the % FIFA composition between net-pen and control sites for each species (Anderson, 2017). As this technique yields pseudo-F statistics, P-values can be calculated using permutations that yield P-values with significance considered for P<0.05. Due to the small and differing sample sizes, we conducted 100,000 permutations to compute a nonparametric PERMANOVA (Steeves et al., 2018). Univariate boxplot comparisons were used to identify which FIFA were driving significant differences in FIFA profiles.

### Detecting Local Changes in Recipient Food Web Biomass Densities

Hydroacoustic surveys were conducted in 2017 to determine the horizontal and vertical spatial distribution of fish surrounding the net-pen facility to detect local aggregations of fish densities surrounding the point-source subsidy input, indicating potential local increases in productivity. One day and one night survey was conducted on July 5 (day survey from 14:30-19:00) and July 6 (night survey from 23:45 - 04:00) 2017 following a 2000 m transects leading away from the net-pen facility (see map inset in Fig. 6). All surveys were conducted following the standard operating procedures outlined in Parker-Stetter et al. (2009) (see Appendix 2 for method details).

To determine if there were significantly higher densities of fish closer to the point-source subsidy input location, Echoview acoustic post-processing software version 7.1.36.30718 was used to convert the cleaned fish count data into fish density values (see Supplementary Methods for details). Fish density values were extracted from Echoview and linear regressions (a = 0.05) were conducted to compare fish density to distance from net-pen for both night and day surveys. Non-linear data was log transformed prior to statistical analysis.

### Detecting Regional Changes in Receiving Food Web Biomass

Fish biomass data from five sites across Georgian Bay (Fig. 7) were obtained from the MNRF’s 2017 Broad-scale Monitoring program (see UGLMU, 2018 for detailed collection methods), to determine if the off-shore net-pen subsidy in Parry Sound drove asymmetrical increases in the cold water, off-shore (pelagic) biomass. Biomass proportions of all species caught in the 2017 BsM sampling was extracted from the MNRF Annual Report of Fisheries Assessment Projects Conducted on Lake Huron, 2017 and sorted into a thermal guild based on their thermal preference (Hasnian et al., 2018). Species with final temperature preferendum (FTP, the temperature that fish gravitate towards when provided with a broad range of temperatures (Hasnain et al., 2018)) <17°C were classified as cold, 17-25°C were classified as cool, and >25°C were classified as warm (Coker et al., 2001). Total proportion of each thermal guild and proportion of each individual species was calculated. Distribution of total biomass across thermal guilds and species was then compared to see if Parry Sound had disproportionately higher biomass in the cold, off-shore thermal guild relative to sites with no net-pen facilities.

### Detecting Changes in Receiving Food Web Structure

To detect alterations in food web structure driven by the point-source subsidy in Parry Sound, we compared the trophic position of the representative food web collected from Parry Sound and control sites in 2016 and 2017 using a carbon and nitrogen two-source stable isotope mixing model (Post, 2002). There is little to no fractionation of δ^13^C and is used to track basal energy sources (i.e., % littoral carbon use), while δ^15^N fractionates from resource to consumer and thus is used to track trophic position. To increase the number of control sites, additional δ^15^N and δ^13^C data was obtained for comparable species from numerous control sites around Lake Huron in 2017 as part of various OMNRF research projects (UGLMU, 2018) (see Appendix 1: Figure S1 for map of sites). All information regarding collection of these samples can be found in the OMNRF Annual Report of Fisheries Assessment Projects Conducted on Lake Huron, 2017 (UGLMU, 2018). Non-lipid corrected C and N values were used as the majority of C:N were low (81% < 3.5 and 92% < 4) (Post et al., 2007).

The following equations were used to calculate trophic position for the mobile, generalist top predators, which incorporate coupling of pelagic and littoral energy channels ((Post, 2002):

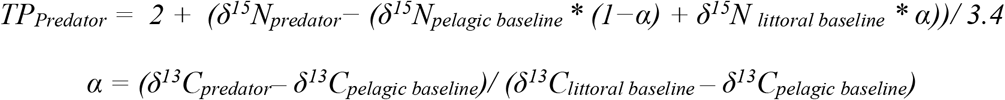

where δ^15^N_predator_ and δ^13^C_predator_, are the nitrogen and carbon stable isotope values for predatory fish collected (lake trout, walleye, smallmouth bass) and δ^15^N_pelagic baseline,_ δ^13^C_pelagic baseline,_, δ^15^N _littoral baseline_, δ^13^C_littoral baseline_ are the nitrogen and carbon stable isotope values for mussels/zooplankton and mayflies respectively. Coupling is incorporated into trophic position calculation through α, which weighs the trophic position estimate according to baseline contributions (Vander Zanden et al., 1999; Vander Zanden et al., 2000).

Trophic position of forage fish was calculated using the following equation, which does not incorporate coupling (Post, 2002):

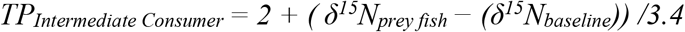

where δ^15^N_prey fish_ and δ^15^N_baseline_ are the nitrogen stable isotope values of forage fish and associated baseline (e.g., littoral forage fish would use littoral baselines). To test for significant differences in trophic position between subsidy and control sites, ANOVA with TukeyHSD post-hoc analysis was conducted (a = 0.05).

All statistical analysis was completed using the R statistical computing package, version 3.6.1 (R Core Team 2019). Full raw data and associated R code will be available on GitHub upon manuscript acceptance.

## Results

### Net-pen Aquaculture Feed Indicator Fatty Acid Selection

PCA of all fatty acids (FA) composing >1% of total FA composition for each forage fish, baseline, and feed sample from net-pen and control sites in 2016 and 2017 successfully separated net-pen feed sources from natural biota across PC1 (Fig 2. (a), (b)). PC1 explained 38.3% of total variation of fatty acid composition in 2016 and 38.8% in 2017. This variance was primarily driven by 3 FA in 2016 (higher % in feed: 18:2n6, 18:1n9, 20:1n9), and 4 FA in 2017 (higher % in feed: 18:2n6, 18:3n3, 18:1n9, 16:1n7), which all loaded significantly onto PC1 and in the direction of the net-pen feed (i.e., PC score > 0.3). These FA were retained as ‘feed indicator fatty acids’ (FIFA) to be used in subsequent analysis comparing individual species FIFA compositions between net-pen and control sites.

### Asymmetrical Uptake of Net-Pen Subsidies Across Thermal Guilds

PCA on FIFA profiles of individual species between net-pen and control sites in 2016 and 2017 show differences in the degree of separation of net-pen and control populations among thermal guild and trophic level. The cold-water top predator (Fig. 4(a; *i*)) and forage fish (Fig. 4(b; *i*)), as well as the cool-water forage fish (Fig. 3(d; *i*); Fig. 4(c; *i*)) demonstrate the highest degree of separation between net-pen and control populations. The cool-water top predator demonstrates some degree of separation between net-pen and control populations (Fig. 3(a; *i*); Fig. 4(c; *i*)), and the warm-water top predator shows some separation of populations in 2016, but a complete overlap with control populations in 2017 (Fig. 3(b; *i*); Fig. 4(d; *i*)). Euclidean distances between cluster centroids of net-pen and control populations demonstrate a negative relationship between species thermal preference (i.e., FTP) and centroid distance, where cold water species tend to have the greatest distance between populations and warm water species the least (Figure 5).

**Figure 3.**
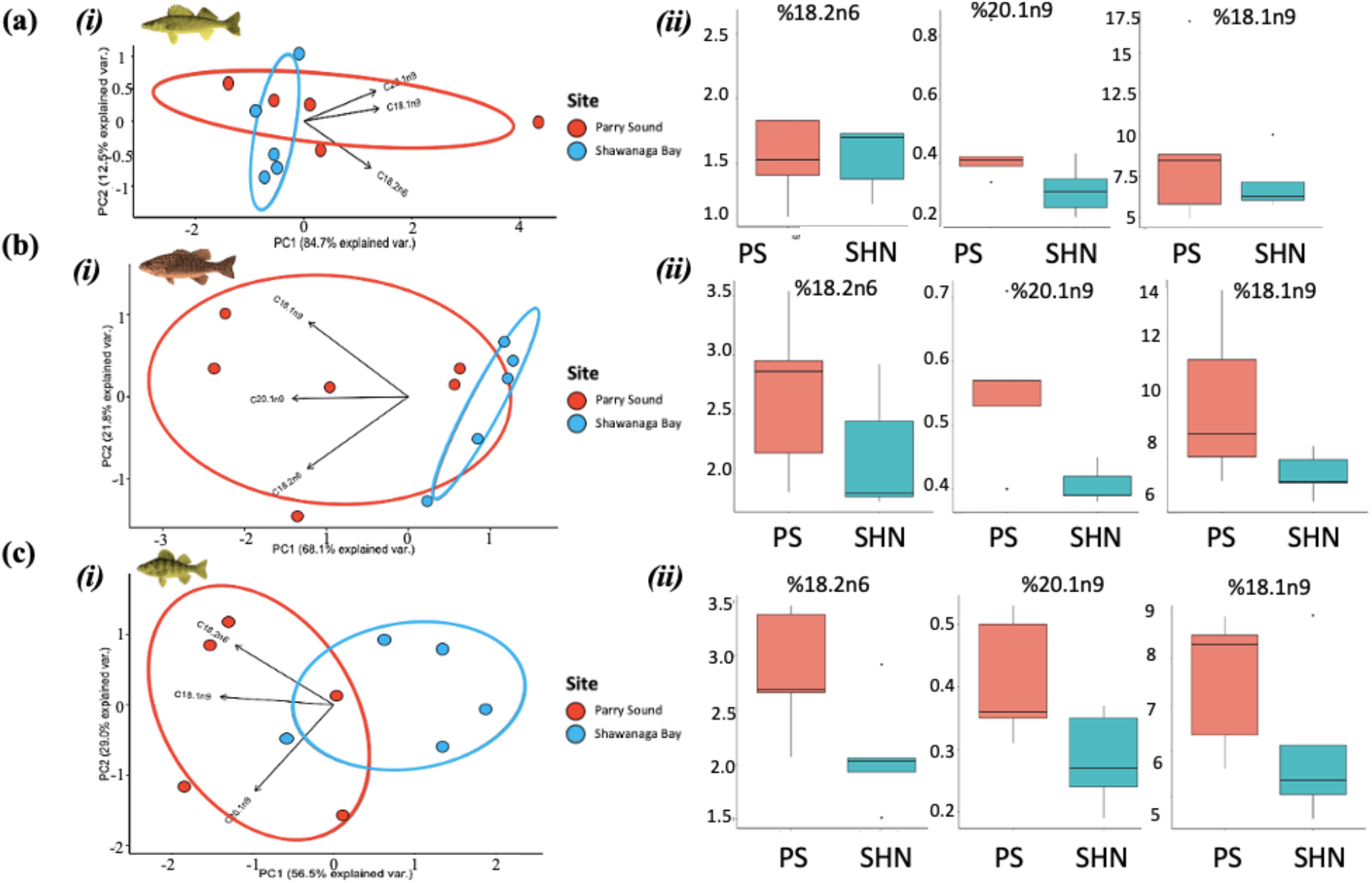
(i) PCA and (ii) univariate boxplot comparisons of feed indicator fatty acids (FIFA) for (a) cool and (b) warm generalist, top predators and (c) cool water forage fish collected from Parry Sound (PS) (anthropogenic subsidy input site) and control sites (Shawanaga Bay, SHN) in 2016.

**Figure 4.**
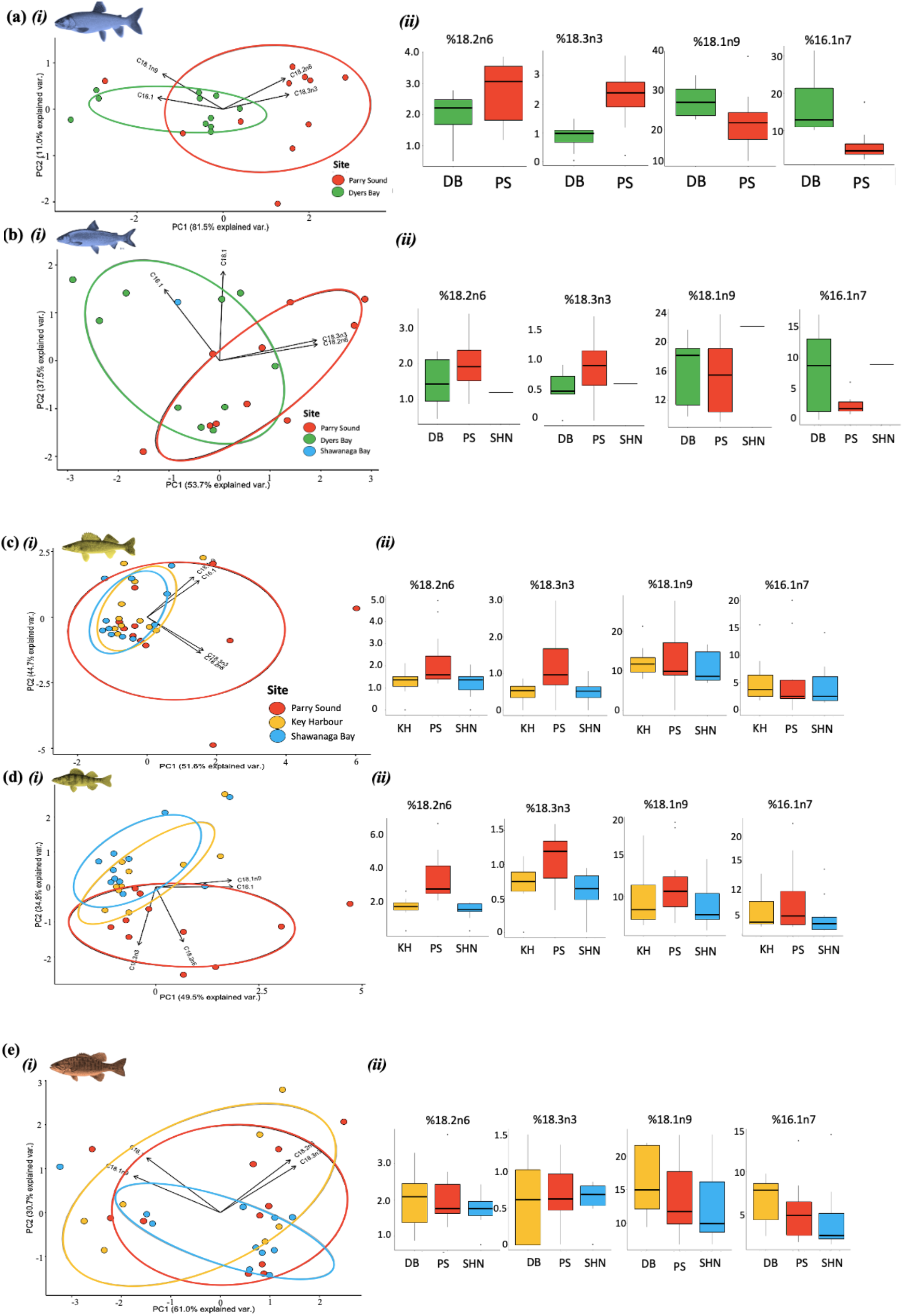
(i) PCA and (ii) univariate boxplot comparisons of feed indicator fatty acids (FIFA) for (a) cold, (c) cool, and (e) warm generalist top predators, and (b) cold and (d) cool water forage fish collected from Parry Sound (PS) (anthropogenic subsidy input site) and control sites (Dyers Bay, DB; Shawanaga Bay, SHN; Key Harbour, KH) in 2017.

**Figure 5.**
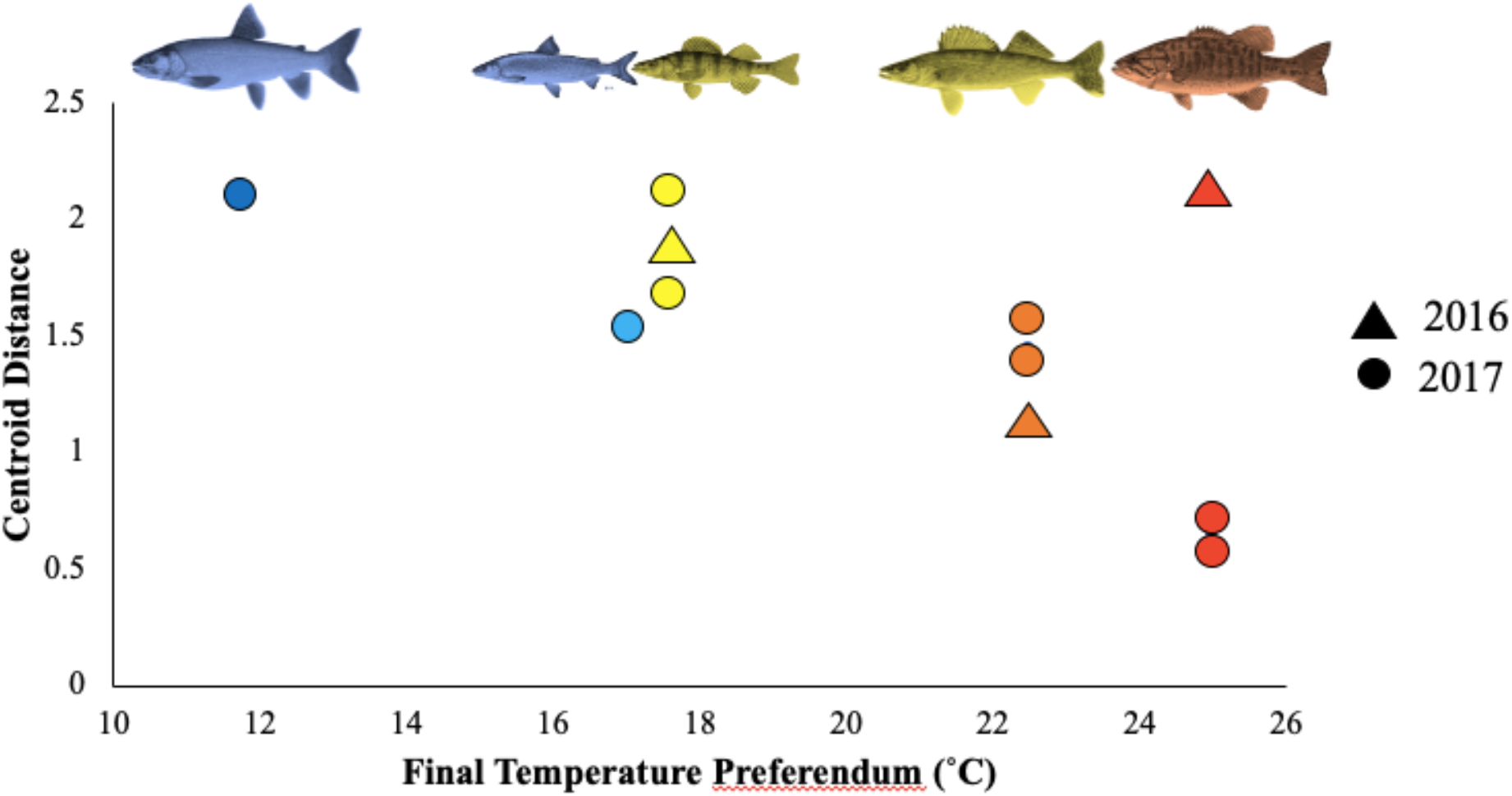
Euclidean distance between centroids of net-pen and control sites, calculated separately for each species and year, in relation to species’ final temperature preference. Blue represents species in the cold thermal guild, yellow and orange are species in the cool thermal guild, and red is species in the warm thermal guild.

**Figure 6.**
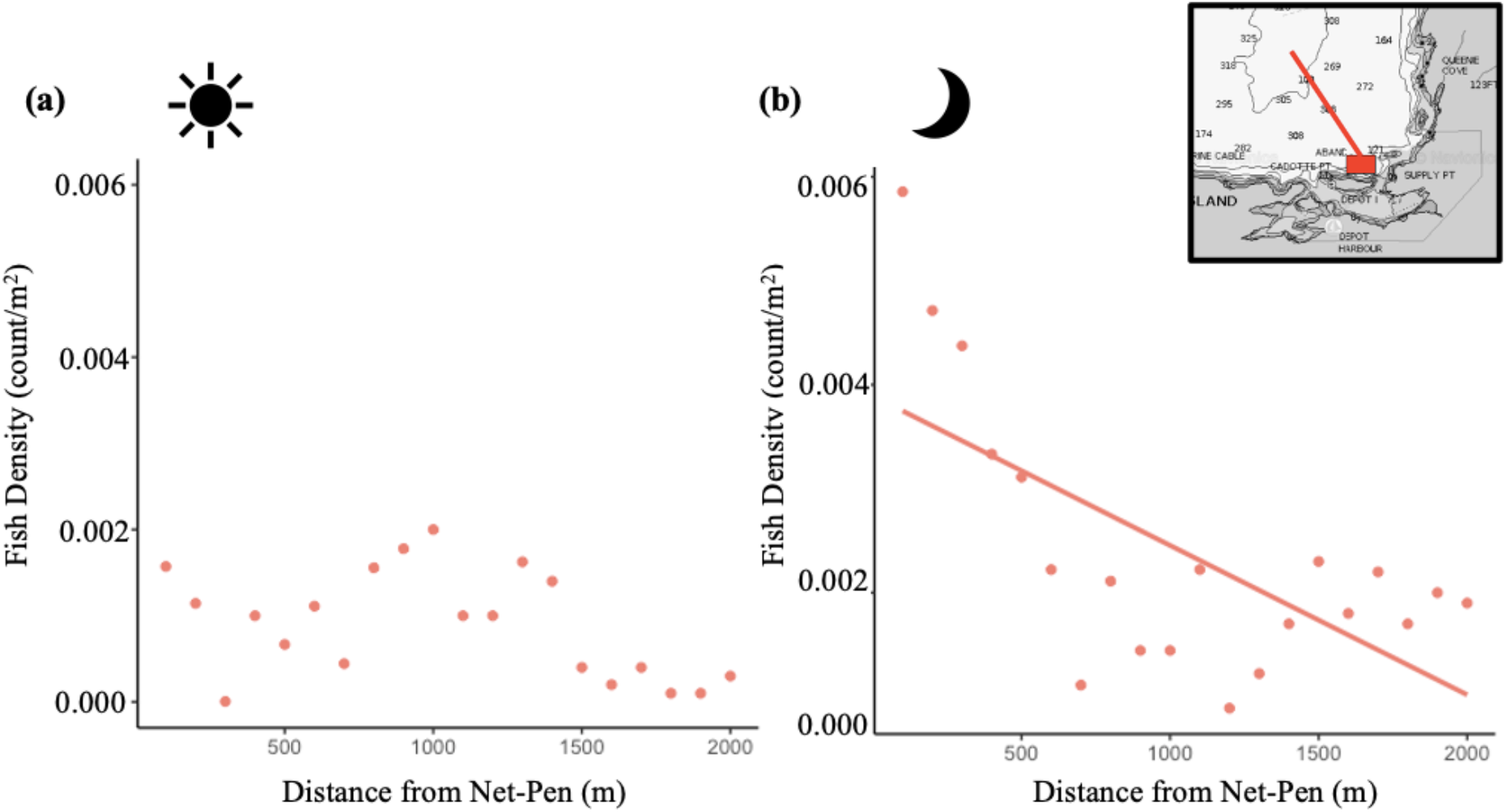
Change in fish density (fish count/m^2^) relative to distance from net-pen facility (m) in Parry Sound for (a) day time transect and (b) night time transect in July 2017. Map inset shows transect location in relation to net-pen facility in Parry Sound (red box), Georgian Bay. Fish density declined with increasing distance from net-pen during the day, however this decline was not significant (R^2^=0.158, p = 0.0821). Fish density significantly declined with increasing distance from net-pen facility at night (R^2^ =0.43, p <0.0001).

**Figure 7.**
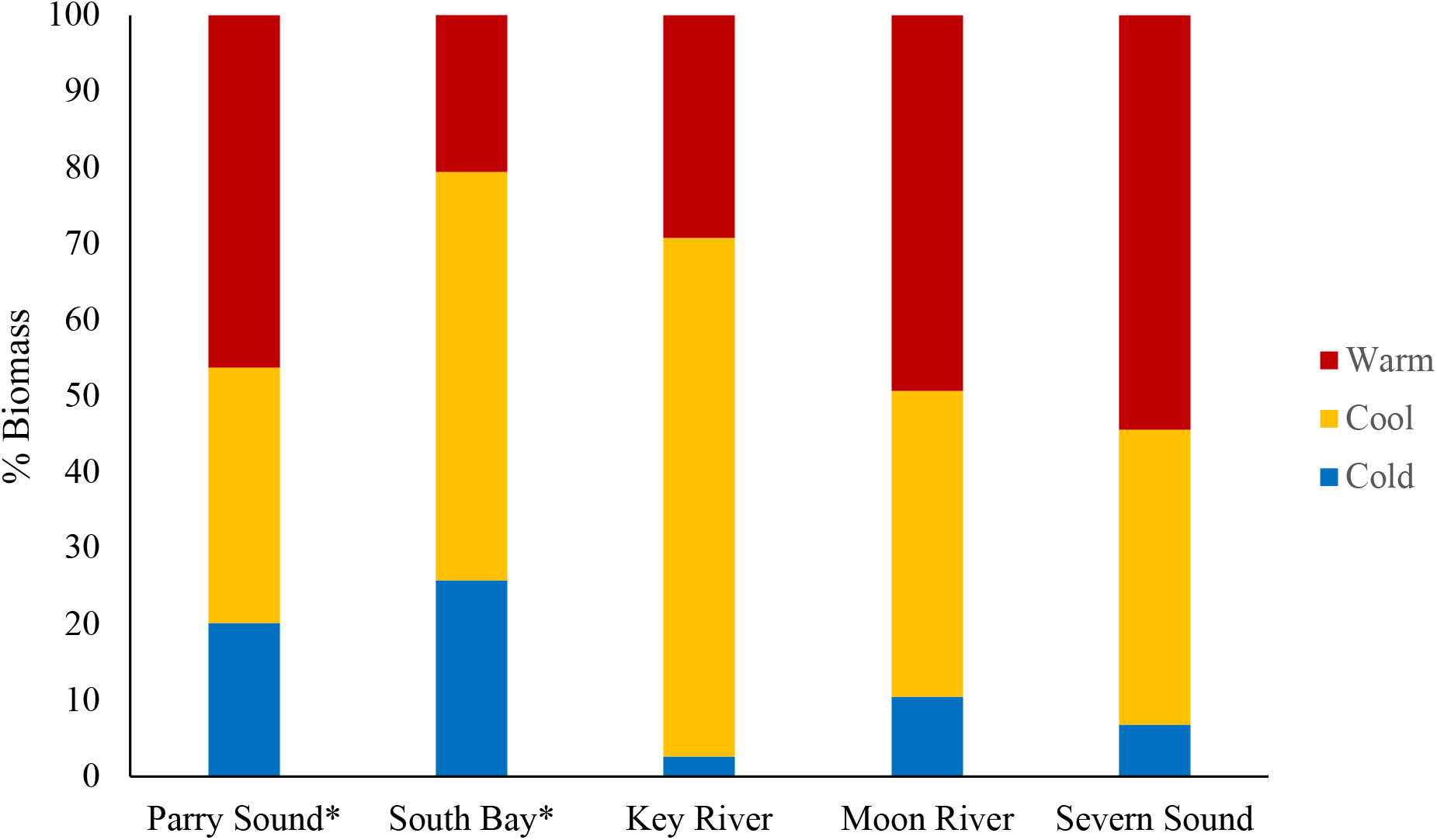
Distribution of biomass proportion across thermal guilds of all species collected from MNRF BSM sampling in Georgian Bay, Lake Huron in 2017. Thermal guilds are indicated by colour, asterisks next to site name indicate sites with presence of net-pen facility. Parry Sound and South Bay, both of which contain net-pen aquaculture facilities, demonstrate higher proportion of species biomass in the cold thermal guild (off-shore species) relative to control sites. Specifically, cold water species biomass in Parry Sound is 2.0 – 7.9 times higher than control sites sampled throughout Georgian Bay, where no net-pen aquaculture facilities were located. High relative biomass of cold water species in Parry Sound to control sites was not reflected in the cool and warm water guilds.

**Figure 8.**
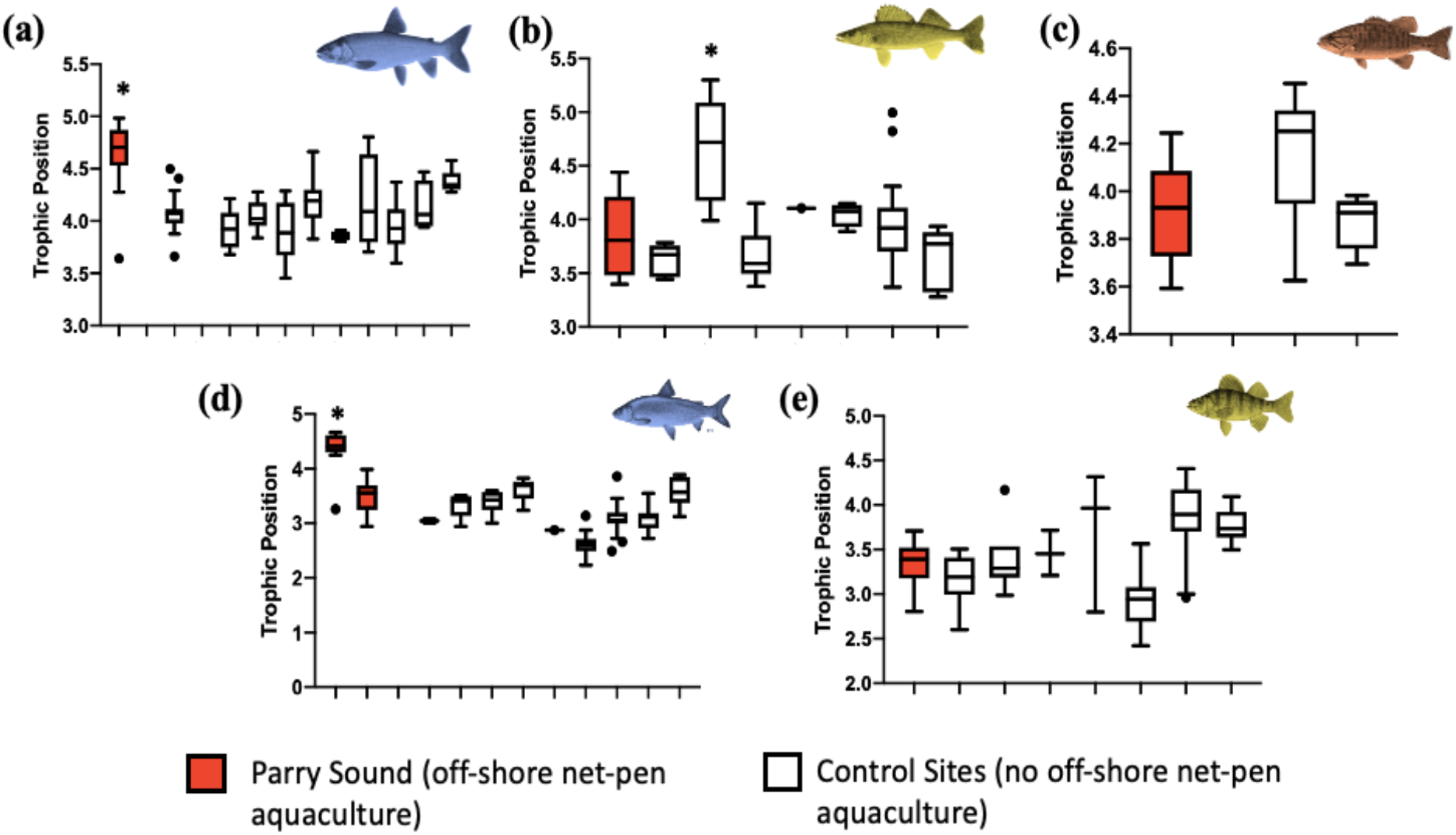
Trophic position of (a) cold, (b) cool and (c) warm generalist top predators, and (d) cold and (e) cool water forage fish collected from Parry Sound (indicated in red) and control sites in 2017. Samples are a combination of those collected by our research team and the MNRF. Asterisks indicate significant differences based on Tukey post-hoc analysis.

PERMANOVA comparing the % FIFA composition between net-pen and control populations of each species shows that cold-water top predators (*p* = 0.0036), cold-water forage fish (*p* = 0.0304) and cool water forage fish (*p* = 0.0343) have the most significantly different FIFA profiles relative to control sites in 2017. The 2016 cool-water forage fish population had the most significant difference in FIFA populations across both years (*p* = 0.0008). In the cold-water top predator and forage fish these significant differences are driven by higher levels of linoleic acid (18:2n6) and linolenic acid (18:3n3) in net-pen populations relative to control (Fig. 4, (a; *ii*); (b; *ii*)). In the cool-water forage fish, significant differences are driven by higher levels of all FIFA in net-pen populations for both 2016 and 2017. The cool-water top predator (i.e., Walleye) net-pen associated FIFA composition was significantly different from control populations in 2017 (*p* = 0.0384) but not in 2016 (*p* = 0.378). Consistent with cold-water species, Walleye also demonstrated higher levels of linoleic acid (18:2n6) and linolenic acid (18:3n3) in net-pen populations relative to control in 2017 (Fig. 4, (c; *ii*)). Despite not demonstrating significant differences in overall FIFA profiles in 2016, univariate boxplots show Walleye still exhibited higher levels of FIFA relative to the control population (Fig. 3, (a; *ii*)), The warm-water top predator (i.e., Smallmouth bass) showed no difference in FIFA profiles in 2017 (*p* = 0.633) and a marginally significant difference in all 3 FIFA in 2016 (*p* = 0.056). Univariate boxplots results show that this marginal significance was driven by higher levels of all 3 FIFA in 2016 net-pen associated smallmouth bass.

### Asymmetric Changes in Recipient Food Web Productivity

Linear regression of fish density surrounding net-pen facility demonstrate increased local aggregations of fish density closer to the net-pen facility than farther away (Figure 6). The strength of aggregation around the net-pen was stronger at night than during the day. While both day and night surveys show decreases in fish density with increasing distance from the net-pen, only the night transect was significant (Night: *p* = 0.0089; Day: *p* = 0.4595). The net-pen is located in 75 m of water and the transect led away from shore, thus we can eliminate the possibility that this effect is caused by a near-shore, littoral increase in fish densities.

Comparison of biomass distribution of cold, cool and warm water species in net-pen (Parry Sound and South Bay) vs. 4 control sites across Georgian Bay, demonstrate Parry Sound and South Bay have higher proportions of cold-water biomass than control sites (Figure 7), which was not reflected in the cool- and warm-water guilds. A large proportion of the cold water biomass in Parry Sound is comprised of Lake Trout (12.46%), Cisco (2.94%) and Alewife (1.21%), which are three cold-water species that were much less abundant throughout the Georgian Bay (see Supplementary Table 3 for % biomass of species at sites across Georgian Bay).

### Asymmetric Changes in Recipient Food Web Structure

The ANOVAs comparing trophic position of cold, cool, and warm water species collected across Lake Huron found that the cold-water top predator and cold water forage fish have significantly higher trophic positions in Parry Sound relative to any other control site in Lake Huron (F = 40.328, *p* = <0.001; F = 48.582, *p* = <0.001, respectively) (Figure 6 (a), (d)). Cool-water top predators and forage fish exhibited significant differences in trophic position between sites across Lake Huron (F = 15.352. *p* = <0.001; F = 18.878, *p* = <0.001, respectively), however this significance was not driven by higher trophic position in Parry Sound relative to all control sites (Figure 6 (b),(e)). Warm-water top predators did not exhibit significant differences between net-pen and control sites (F=2.948, p = 0.06219) (Fig. 6(c).) It should be noted that the MNRF data only included pelagic baselines, therefore coupling was not possible to calculate and incorporate into trophic position calculation of MNRF sampled generalist predators. Statistical outcomes were consistent regardless of if coupling was incorporated or not, thus coupling was incorporated where possible to provide more accurate trophic position estimates when able.

## Discussion

Despite the fact that accessibility may fundamentally alter the influence of a subsidy on resultant food web dynamics, surprisingly this aspect of subsidies remains relatively unexplored. Here, we use a natural experiment to determine how species traits, particularly thermal tolerance, influenced accessibility to an off-shore point-source anthropogenic subsidy, and in turn show that differential accessibility drives asymmetric changes in food web productivity and structure. Our results suggest that species traits can produce a gradient in subsidy accessibility, as cold-water adapted species show the greatest evidence of subsidy consumption, and warm-water species demonstrate little to none. Notably, our feeding results also scale up to show consistent asymmetric changes in productivity and food web structure such that cold-water species, for example, consume most of the subsidy and have elevated biomass and trophic position.

Recent work has argued that climate change can be expected to generally have asymmetrical impacts on habitat (e.g., habitats differentially warmed by climate change) that lead to organismal behaviour that asymmetrically rewires whole carbon pathways. Bartley et al (2018) argued that differential habitat changes and species behaviour (traits) then drive wholesale food web rewiring. Our results here suggest that such impacts of climate change rewiring may extend more broadly to other aspects of global change. Specifically, here we showed that the placement of net-pen aquaculture in cold-water habitats (i.e., a global change that is necessarily habitat dependent and so asymmetrical) produces stronger interactions as a function of species traits (i.e., thermal tolerance) and so functionally rewires this ecosystem, altering food web structure and productivity with yet unknown stability implications (Blanchard et al., 2015; Bartley et al. 2018). Like climate change, net-pen inputs can differentially alter macrohabitats (littoral versus pelagic) and so drive asymmetric rewiring. It remains to be seen whether human impacts that cross multiple habitats generally have asymmetrical impacts, and if they do, this suggests a general rewiring result of global change on spatial food webs.

The use of fatty acids to trace specific prey items through food webs has provided a powerful tool for understanding how subsidies move through and are utilized by the surrounding ecosystem. Fatty acid composition analysis has been widely used in the aquaculture literature to identify the presence of net-pen feed in the diet of surrounding fish species in both marine and freshwater systems (Kullman et al., 2009; Fernandez-Jover et al., 2011; Johnson et al., 2018). The presence of marine and terrestrial derived protein and oil in fish feed differentiate the fatty acid composition of the feed from natural food sources (Fernandez-Jover et al., 2011). Fatty acids are highly conserved throughout the food web, which provides the ability to trace net-pen feed through a surrounding food web because differences in fatty acid composition between net-pen and control populations is proportional to the amount of feed consumed (Fernandez-Jover et al., 2011).

We demonstrated how subsidy accessibility, measured through amount of consumption of net-pen feed, depends on species’ traits (e.g., thermal tolerance). Cold-water mobile generalists and forage fish had significant differences in feed indicator fatty acid (FIFA) composition relative to control populations (little overlap in FIFA clusters, large centroid distance (Fig. 4; Fig. 5)). This suggests their diet is significantly different from control diets due to the presence of feed. Cold-water forage fish were less significantly different than the mobile generalist, which may be a factor of the different dominant pelagic forage fish found in each locations (Alewife in Parry Sound, Rainbow Smelt in Dyer’s Bay). In our study, cool-water forage fish had the most significant difference in FIFA profile in 2016, and while still significant in 2017, not as significant as the cold-water generalist and forage fish. Cool-water mobile generalists’ FIFA profiles were also significantly different from control populations, however the signal was not as strong (centroid distance not as far, more cluster overlap) as in the cold-water species. The cool-water generalist and forage fish also displayed higher variation in the Parry Sound populations relative to controls and cold-water species. This suggests there may be more individual variation in the cool-water guild, where some individuals strongly incorporate FIFA and some were no different than other control fish. Warm-water mobile generalists demonstrated high overlap between net pen and control sites, suggesting little to no feed uptake and low access to net-pen subsidies in 2017. Marginally significant differences between net pen and control sites in 2016 indicate some individuals may have accessed net-pen subsidies that year. Some evidence of access in 2016 may be due to warmer temperatures that summer (Parry Sound Historical Weather Data, Environment Canada), which may have made a larger warmer zone and thus able to forage farther and potentially access net-pen resources. It should be noted that littoral baselines (i.e., mayflies) had similar values of linolenic acid (18:3n3) and higher values of palmitoleic acid (16:1n7) relative to the feed due to the input of terrestrial oils in the feed (Appendix 1: Figure S1). This may drive similarities in FIFA profiles of littoral cool and warm water species; therefore, it is important to focus on fatty acids that are significantly different from the feed as stronger feed indicators in these species. Here, our results provide evidence of a gradient in net-pen feed uptake, that is correlated with the gradient in thermal preferences, suggesting thermal tolerances limit the ability of surrounding species to access point-source subsidies. Further, our results suggest the intriguing possibility that the population of cold-water species (e.g. lake trout) generally respond to consume from the cold-water feed site, while more thermally generalized species (cool-water) show greater variation in response suggesting individual variation in amount of feed intake.

By looking at changes in local and regional fish density and biomass patterns, we were able to simultaneously detect asymmetrical increases in biomass of the surrounding food web along the accessibility gradient. These changes in biomass suggest asymmetrical access to net-pen resources may drive differential productivity in the surrounding food web (e.g., cold-water species elevated). Investigation into patterns of fish density surrounding the net-pen showed increased density locally around the net-pen, suggesting surrounding species aggregate to forage on the released net-pen subsidy. These aggregations were also stronger at night, when fish are actively foraging. Our results follow previous aquaculture literature that demonstrates increased fish densities surrounding fish farm operations in both marine and freshwater systems (Ferndandez-Jover et al., 2007; Johnston et al., 2010; Rennie et al., 2019) Investigation of the distribution of regional fish biomass demonstrated that Parry Sound had a higher proportion of biomass of cold-water species relative to sites with no point-source subsidy present. The high biomass of cold-water species was largely driven by higher proportion of biomass of lake trout relative to all other sites studied in Georgian Bay, Lake Huron. Parry Sound also supports a natural reproducing lake trout population that does not undergo yearly stocking (UGLMU, 2018). South Bay was the only site that had comparable proportion of total biomass in the cold-water guild. However, here the guild is dominated by whitefish and there are no lake trout present, it also supports a lower diversity of cold-water fish than Parry Sound (Appendix 1: Table S3). The significantly higher proportion of offshore biomass in Parry Sound, and presence of a naturally reproducing off-shore top predator that strongly accesses net-pen feed, suggests net-pen subsidies may be increasing off-shore productivity. Further research investigating how these released subsidies are influencing basal food web productivity (i.e., zooplankton biomass) will be helpful in elucidating the mechanism through which net-pen subsidies are driving significantly higher cold-water guild biomass in Parry Sound.

Through examination of trophic position of net-pen associated species and control species throughout Lake Huron, we detected significant increases in trophic position of cold-water species in Parry Sound relative to all other sites in Lake Huron, which was not reflected in cool and warm water species. As higher trophic positions indicate longer food chains (Vander Zanden et al., 1999), these results suggest that thermal guilds with the greatest accessibility to net-pen subsidies (in this case cold-water guild) have a significantly longer food chain than guilds with less access. Therefore, the gradient in subsidy accessibility drives asymmetric changes in surrounding food web structure (i.e., lengthening of cold water food chain, no lengthening of cool-warm water food chain). These results are supported by previous work conducted by Johnson et al. (2018) that provided the first evidence of higher trophic position in cold-water species in net-pen associated sites. Asymmetrical changes in food web structure can fundamentally rewire whole food webs (Bartley et al., 2019), which influences whole food web stability and function in yet unknown ways. As accessibility is a main driver of these asymmetric changes in surrounding food web that may influence whole food web stability, our research highlights the importance of looking at subsidy accessibility when trying to understand or predict subsidy-driven food web dynamics.

Here, we provide the first evidence that accessibility gradients to subsidies do exist and can largely influence the strength of subsidy assimilation and changes in resultant food web productivity and structure. To determine if subsidy accessibility is ubiquitous across ecosystems (i.e., is this a characteristic of subsidies that is always relevant), more empirical studies investigating this concept are needed. Additionally, theoretical work focused on understanding how asymmetrical changes in productivity and structure based on differential accessibility scenarios influence food web stability and function are needed to understand the dynamical implications of subsidy accessibility.

Global change and increasing human populations are continually altering the flow of natural and anthropogenic subsidies throughout ecosystems, thus it is imperative to understand how these subsidies, in combination with global changes, will influence future food web stability and function. Elucidating key subsidy characteristics that drive changes in recipient food web structure and function is essential. Here, we contribute another key subsidy characteristic, subsidy accessibility, that has yet to be considered in subsidy literature. While this study shows how accessibility is key in determining food web responses to an anthropogenic subsidy, we argue this concept can be more widely applied to all subsidies and is a key characteristic of subsidy-receiving food web interactions that must be considered when trying to understand subsidy impacts on receiving ecosystem stability and function under continued global change.

## Acknowledgments

This project was in part funded by the University of Guelph’s Canada First Research Excellence Fund project “Food from Thought,” awarded to K.S.M, and from Aqua-Cage Fisheries Ltd. We would like to thank the Ontario Ministry of Natural Resources and Forestry (OMNRF) and their multiple sampling programs that contributed to this study.

## Literature Cited

Anderson, M. J. 2017. Permutational Multivariate Analysis of Variance (PERMANOVA). Wiley StatsRef: Statistics Reference Online:1–15.

Bartley, T., K. S. Mccann, and C. Bieg. 2018. Food web rewiring in a changing world.

Blanchard, J. L. A. A rewired food web. Nature 527, 7–8 (2015).

Coker, G. A., C. B. Portt, and C. K. Minns. 2001. Morphological and Ecological Characteristics of Canadian Freshwater Fishes. Canadian Manuscript Report of Fisheries and Aquatic Sciences 2554:iv+89.

DeBruyn, A. M. H., D. J. Marcogliese, and J. B. Rasmussen. 2003. The role of sewage in a large river food web. Canadian Journal of Fisheries and Aquatic Sciences 60:1332–1344.

Debruyn, A. M. H., K. S. Mccann, J. B. Rasmussen, A. M. H. Debruyn, K. S. Mccann, and J. B. Rasmussen. 2020. Migration Supports Uneven Consumer Control in a Sewage-Enriched River Food Web Published by: British Ecological Society Linked references are available on JSTOR for this article: Migration supports uneven consumer control in a sewage-enriched river food 73:737–746.

Eveleigh, E. S., K. S. McCann, P. C. McCarthy, S. J. Pollock, C. J. Lucarotti, B. Morin, G. a. McDougall, D. B. Strongman, J. T. Huber, J. Umbanhowar, and L. D. B. Faria. 2007. Fluctuations in density of an outbreak species drive diversity cascades in food webs. Proceedings of the National Academy of Sciences 104:16976–16981.

Fernandez-Jover, D., P. Sanchez-Jerez, J. T. Bayle-Sempere, C. Valle, and S. Derrien. 2008. Seasonal patterns and diets of wild fish assemblages associated with Mediterranean coastal fish farms. ICES Journal of Marine Science 65:1153–1160.

Hasnain, S. S., M. D. Escobar, and B. J. Shuter. 2018. Estimating thermal response metrics for North American freshwater fish using Bayesian phylogenetic regression. Canadian Journal of Fisheries and Aquatic Sciences 75:1878–1885.

Huxel, G. R., K. McCann, and G. a. Polis. 2002. Effects of partitioning allochthonous and autochthonous resources on food web stability. Ecological Research 17:419–432.

Johnson, L. E., B. McMeans, N. Rooney, M. Gutgesell, R. Moccia, and K. S. McCann. 2018. Asymmetric assimilation of an anthropogenic resource subsidy in a freshwater food web. Food Webs 15:e00084.

Lee, K. Y., L. Graham, D. E. Spooner, and M. a. Xenopoulos. 2018. Tracing anthropogenic inputs in stream foods webs with stable carbon and nitrogen isotope systematics along an agricultural gradient. PLoS ONE 13:1–19.

Leroux, S. J., and M. Loreau. 2008. Subsidy hypothesis and strength of trophic cascades across ecosystems. Ecology Letters 11:1147–1156.

McCann, K. S. 2005. The dynamics of spatially coupled food webs:513–523.

Nakano, S., and M. Murakami. 2001. Reciprocal subsidies: Dynamic interdependence between terrestrial and aquatic food webs. Proceedings of the National Academy of Sciences of the United States of America 98:166–170.

Newsome, T. M., J.a. Dellinger, C. R. Pavey, W. J. Ripple, C. R. Shores, A. J. Wirsing, and C. R. Dickman. 2015. The ecological effects of providing resource subsidies to predators. Global Ecology and Biogeography 24:1–11.

Parker-Stetter, S. L., L. G. Rudstam, P. J. Sullivan, and David M. Warner. 2009. Standard operating procedures for fisheries acoustic surveys in the Great Lakes. Special Publication 09-01 January 2009. Great Lakes Fishery Commission.

Polis, G. a., W. B. Anderson, and R. D. Holt. 1997. Toward an integration of landscape and food web ecology: The dynamics of spatially subsidized food webs. Annual Review of Ecology and Systematics 28:289–316.

Polis, G. a., and D. R. Strong. 1996. Food web complexity and community dynamics. American Naturalist 147:813–846.

Post, D. M. 2002. Using stable isotopes to estimate trophic position: models, methos, and assumptions. . Ecology 83:703–718.

Post, D. M., C. A. Layman, D. A. Arrington, G. Takimoto, J. Quattrochi, and C. G. Montaña. 2007. Getting to the fat of the matter: Models, methods and assumptions for dealing with lipids in stable isotope analyses. Oecologia 152:179–189.

Rennie, M. D., P. J. Kennedy, K. H. Mills, C. L. Podemski, C. M. C. Rodgers, C. Charles, L. E. Hrenchuk, S. Chalanchuk, P. J. Blanchfield, and M. J. Paterson. 2019. Impacts of freshwater aquaculture on fish communities: A whole-- ecosystem experimental approach:870–885.

Rodewald, A. D., L. J. Kearns, and D. P. Shustack. 2011. Anthropogenic resource subsidies decouple predator-prey relationships. Ecological Applications 21:936–943.

Sears, A.., R. D. Holt, and G. a. Polis. 2004. Feast and Famine in Food Webs: The Effects of Pulsed Productivity. Pages 359–386 *in* G.A. Polis, G.R. Huxel and M. Power, eds. Food Webs at the Landscape Scale: The Ecology of Trophic Flow across Habitats. University of Chicago Press, Chicago, IL.

Steeves, H. N., B. McMeans, C. Field, C. Stewart, M. T. Arts, A. T. Fisk, C. Lydersen, K. M. Kovacs, and M. A. Macneil. 2018. Non-parametric analysis of the spatio-temporal variability in the fatty-acid profiles among Greenland sharks. Journal of the Marine Biological Association of the United Kingdom 98:627–633.

Takimoto, G., T. Iwata, and M. Murakami. 2002. Seasonal subsidy stabilizes food web dynamics: Balance in a heterogeneous landscape. Ecological Research 17:433–439.

Upper Great Lakes Management Unit (UGLMU). (2018). Annual Report of Fisheries Assessment Projects Conducted on Lake Huron, 2017. Ministry of Natural Resources and Forestry, Report PS-LHA-2017-01.

Vander Zanden, M. J., and J. B. Rasmussen. 1999. Primary Consumer 13 C and 15 N and the Trophic Position of Aquatic Consumers. Ecology 80:1395–1404.

Vander Zanden, M. J., B. J. Shuter, N. Lester, and J. B. Rasmussen. 1999. Patterns of food chain length in lakes: A stable isotope study. American Naturalist 154:406–416.

