## Appendix S2 for "Subsidy Accessibility Drives Asymmetric Food Web Responses"

Ecology

### Appendix S2: Detailed Hydroacoustic Data Collection Methods

*Transect set up and BioSonics hydroacoustic data collection*

In July 2017, both night-time and day-time SONAR surveys were conducted in Lake Huron along one transect starting at, and leading away from the Aqua-Cage Fisheries cage-culture (Parry Sound, Ontario, Canada; Fig. 6). The day-time survey was conducted from 2:30pm to 7pm, while the night-time survey was conducted from 11:45pm to 4 am. The thermocline was estimated based on temperature profiles, sitting at approximately 10m below the surface.

Hydroacoustic procedures were based on Parker-Stetter et al.'s (2009) "Standard operating procedures for fisheries acoustic surveys in the great lakes". Acoustic data was collected with a BioSonics DT-X extreme autonomous portable scientific echosounder equipped with a 430 kHz and a 120 kHz elliptical split-beam transducer, calibrated by the standard sphere method (Foote et al. 1987). For the purpose of this study, only the 120 kHz frequency echogram returns were analysed due to target specimen size (fish as opposed to zooplankton). The transducer was deployed off the stern of the vessel at a depth of 1m where it was dragged along the transect at a survey speed of 5.5-6 km/h. Ping rates of 0.8 pings/s were used with a pulse duration of 0.5 ms to allow for the discrimination of fish from the bottom, avoiding 'shadow bottom'. Acoustic signals were collected with BioSonics Visual Acquisition Software (version

4.1), and output files were stored on a laptop computer hard drive. Vessel position was integrated into the BioSonics output files by associating each ping return with GPS coordinates.

*Echoview data clean up*

Acoustic echogram files were processed using Echoview acoustic postprocessing software (Fig. 2; version 7.1.36.30718, SonarData). At each transect water temperature and depth were recorded, however, salinity was not measured, therefore this value was not incorporated for calculations of sound speed and absorption coefficient. The calibration values within the Echoview software were compared to that of the calibration settings of the DTX BioSonics Echosounder during the sampling period, ensuring a consistent offset value of 0.4. A surface exclusion zone was determined at a depth of 1m and all data above this line was excluded to avoid any trawling noise pulse manipulation. The best bottom candidate algorithm was used to define the lake bottom due to variation in depth profile. After defining the lake bottom, a linear offset line was added 1m above the bottom line marking the bottom dead zone, in which fish and any other minute biotic and abiotic pulse returns against the bottom of the lake were excluded from analysis.

Background noise removal was conducted by applying bad data regions and by running a background noise removal algorithm. Echoview considers bad data regions as no data which consists of data points which are off transect, below the target layer, or have been subjected to bad weather, interference, ghost bottoms, and echosounder malfunction. Empty water is also removed by applying bad data regions, which excludes volumes of water devoid of targets. The background noise removal scrutinizes the data for acoustic, electrical, and trawl noise by estimating the background noise value for each ping and then subtracting it from the ping's samples. The values used in this algorithm were based on DeRobertis & Higginbottom (2007) and are available in Table 1.

The Method 2 split-beam single target detection algorithm was then applied to isolate single-fish echoes by utilizing aspects and characteristics of the shape of the return pulse. Values from Hrabik et al. (2006) were applied (Table 2). This algorithm allows an echo to be classified as a single target if it meets the following criteria: (1) the echo TS value is a local maximum (larger than surrounding digital samples); (2) the echo TS exceeds a 55-dB threshold; (3) the beam compensation value for the echo is less than 6 dB; (4) the echo pulse duration, which is measured 6 dB from the echo envelope peak, has to fall between 0.8 and 1.5 times the emitted pulse duration; and (5) the standard deviation of all samples within the pulse envelope have to be less than 1.5 (Stockwell et al., 2007).

*Fish count and density calculation*

The previous data clean up procedures yield an echogram that allows for accurate fish count determination along the transect (Fig. 2). These fish counts can thereafter be converted into density values to provide a representation of fish aggregation in relation to the high nutrient densities surrounding the net pen aquaculture. Fish count was separated into bins to avoid any pulse return bias, as pulses are amplified with increased depth. The vertical bin size was based on the value suggested by Parker Stetter et al. (2009) for Lake Huron, at a length of ten meters. The horizontal bin size, however, was altered from Parker Stetter et al.'s (2009) suggestion to allow for the visualization of small-scale changes along the 2000m transect, applying bins of 100 meters instead of 1000m. The fish counts were calculated for each bin, summed within 100m horizontal increments, and divided by the total vertical area of the analysed bin to provide numerical fish densities (fish count/$m^2$) for every 100m along the transect. The counts were also summed by depth layer (horizontally) to provide information on percent fish distribution at differing depths along the transect.

*Statistical analysis*

To determine whether a significant relationship was observed between fish density and transect distance, linear regressions plotting fish density against transect distance were performed in RStudio for the five night transects and the five day transects. Density measurements were plotted for every 100m of transect length for each ~2000m transect, and data yielding non-linear patterns were log transformed. Significance was determined by comparing p-values to an alpha value of 0.05.

**Literature Cited**

DeRobertis, A., and I. Higginbottom. 2007. A post-processing technique to estimate the signal-to-noise ratio and remove echosounder background noise. ICES J. Mar. Sci. 64: 1282-1291.

Foote, K.G., Knudsen, H.P., Vestnes, G., MacLennan, D.N., and Simmonds, E.J. 1987. Calibration of acoustic instruments for fish density estimation: a practical guide. ICES Cooperative Res. Rep. No. 144.

Hrabik, T., Schreiner, D., Balge, M., & Geving, S. (2006). Development of a hydroacoustic survey design to quantify prey fish abundance in the Minnesota waters of Lake Superior. *Minnesota Department of Natural Resources, Investigational Report*, **530**.

Parker-Stetter, S. L., L. G. Rudstam, P. J. Sullivan, and David M. Warner. 2009. Standard operating procedures for fisheries acoustic surveys in the Great Lakes. Special Publication 09-01 January 2009. Great Lakes Fishery Commission.

Stockwell, J. D., Yule, D. L., Hrabik, T. R., Adams, J. V., Gorman, O. T., & Holbrook, B. V. (2007). Vertical distribution of fish biomass in Lake Superior: implications for day bottom trawl surveys. *North American Journal of Fisheries Management*, **27**(3), 735-749.
