## Appendix S1 for "Subsidy Accessibility Drives Asymmetric Food Web Responses"

Ecology

**<u>Appendix 1: Supplementary Tables & Figures</u>**

**Table S1** Composition of feed pellets used by AquaCage Fisheries Inc. in 2016 and 2017. In 2016, only one size of feed was sampled.

| Year | Feed Size (mm) | Fish Size (g) | Protein (min) | Oil (min) | Moisture (max) | Fibre (max) | Ash (max) | Phos (max) | DE (MJ/KG) |
|---|---|---|---|---|---|---|---|---|---|
| 2016 | 6 | 500-1000 | 46% | 24% | 8% | 2% | 10% | 1% | 19.5 |
| 2017 | 4 | 100-400 | 45% | 28% | 8.50% | 1.50% | 12% | NA | 20.2 |
| 2017 | 6 | 400-1000 | 43% | 31% | 8.50% | 1.50% | 12% | NA | 20.2 |
| 2017 | 7.5 | 1000+ | 39% | 32% | 8.50% | 1.50% | 12% | NA | 20.2 |

**Table S2.** Proportion (%) of fatty acids (>1% in system) in feed in 2016 and 2017. Values for 2017 were calculated as an average of fatty acid % in all three feed sizes. Fatty Acid Type describes how fatty acids were characterized in our study: Dietary Fatty Acids (DFA); and Feed Indicator Fatty Acids (FIFA). The year following FIFA indicates which year fatty was a feed indicator. % FIFA are indicated in bold.

| Fatty Acid | Fatty Acid Type | 2016 | 2017 |
|---|---|---|---|
| C14:0 | EDFA | 3.289 | 2.587 |
| C16:0 | EDFA | 19.919 | 19.016 |
| **C16:1n7** | **FIFA_2017** | 5.230 | **9.497** |
| C18:0 | EDFA | 4.587 | 5.818 |
| **C18.1n9** | **FIFA_2016/7** | **23.993** | **30.430** |
| C:18:1n7 | EDFA | 2.073 | unk |
| **C18.2n6** | **FIFA_2016/7** | **17.265** | **22.180** |
| **C20.1n9** | **FIFA_2016** | **3.775** | 0.000 |
| **C18:3n3** | **FIFA_2017** | 1.550 | **5.529** |
| C20:4n6 | EDFA | 0.558 | 0.247 |
| C20:5n3c | EDFA | 3.705 | 2.384 |
| C22:5n3 | EDFA | 0.514 | unk |
| C22:6n3c | EDFA | 3.645 | 0.216 |

**Table S3.** Final temperature preferendum (FTP), thermal guild, and proportion (%) biomass of each species caught in 5 locations as part of the OMNRF 2017 Broad-scale Monitoring Program in Lake Huron. Values were extracted from UGLMU, 2018 Figure 4 (Key River); Fig. 7 (Moon River); Fig. 10 (Parry Sound); Fig 12 (Severn Sound); and Fig. 14 (South Bay). The Broadscale Fish Community Monitoring design used both large mesh and small mesh gillnets. The large mesh gillnets had eight panels (mesh sizes 38 mm to 127 mm stretched mesh) and a total length of 24.85 m. The small mesh gillnets had five panels (mesh sizes from 13 mm to 38 mm stretched mesh) with a total length of 12.5 m. Nets were set overnight and in pairs. Sites were selected using a depth-stratified randomized design using ArcGIS.

| **Species** | **FTP (˚C)** | **Thermal Guild** | **Key River** | **Moon River** | **Parry Sound** | **Severn Sound** | **South Bay** |
|---|---|---|---|---|---|---|---|
| *Alewife* | 16.9 | Cold | 0.10 | 0.10 | 1.20 | 0.34 | 0.34 |
| *Black Crappie* | 23.4 | Cool | 0.26 | 0.10 | - | 0.68 | - |
| *Bluntnose Minnow* | 24.1 | Cool | 0.10 | 0.10 | 0.34 | - | 0.34 |
| *Blackchin Shiner* | 21.8 | Cool | 0.10 | 0.10 | - | - | - |
| *Blacknose Shiner* | | Cool | 0.10 | - | - | - | - |
| *Bowfin* | 30.3 | Warm | 3.07 | 1.17 | 3.59 | 0.68 | 4.47 |
| *Brown Bullhead* | 26.2 | Warm | 6.14 | 7.49 | 2.39 | 2.38 | 3.44 |
| *Bluegill* | 30.2 | Warm | - | - | - | 0.34 | - |
| *Brook Stickleback* | 21.3 | Cool | - | - | - | - | 0.34 |
| *Burbot* | 13.2 | Cold | 1.02 | 6.79 | - | 1.02 | 5.84 |
| *Channel Catfish* | 27.3 | Warm | 0.26 | 1.40 | - | - | - |

| | | | | | | | |
|---|---|---|---|---|---|---|---|
| *Common Shiner* | 21.9 | Cool | - | 0.10 | 0.17 | - | - |
| *Chinook Salmon* | 13.8 | Cold | - | - | - | 3.40 | - |
| *Common Carp* | 27.7 | Warm | - | - | 2.56 | 3.40 | 3.09 |
| *Golden Shiner* | 21.8 | Cool | 0.10 | 0.10 | - | - | - |
| *Emerald Shiner* | 19.3 | Cool | - | - | 0.17 | - | - |
| *Johnny Darter* | 22.8 | Cool | 0.10 | 0.10 | - | - | - |
| *Lake Chub* | 27 | Warm | 0.26 | 0.10 | - | - | - |
| *Lake Herring, Cisco* | 12.4 | Cold | 0.26 | 0.10 | 2.91 | 0.34 | 1.72 |
| *Lake Trout* | 11.8 | Cold | - | 3.04 | 12.31 | - | - |
| *Lake Whitefish* | 12.7 | Cold | - | - | 2.22 | 0.34 | 16.49 |
| *Largemouth Bass* | 28.6 | Warm | 0.10 | 0.10 | 1.54 | 2.72 | - |
| *Longnose Gar* | 27.4 | Warm | 1.54 | 10.53 | 3.93 | 24.15 | - |
| *Logperch* | | Cool | 0.10 | - | - | - | - |
| *Longnose Sucker* | 11.1 | Cold | 0.77 | 0.23 | - | - | - |
| *Mimic Shiner* | | Cool | - | 0.10 | - | 0.34 | 0.34 |
| *Muskellunge* | 25.4 | Warm | - | - | - | 1.70 | - |
| *Northern Pike* | 20.7 | Cool | 20.98 | 21.30 | 9.74 | 7.82 | - |
| *Pumpkinseed* | 27.7 | Warm | 0.26 | 0.47 | 0.17 | 0.68 | - |
| *Rainbow Smelt* | 11.2 | Cold | 0.26 | 0.23 | 0.17 | 0.68 | 1.37 |
| *Rainbow Trout* | 15.5 | Cold | - | - | 0.85 | - | - |
| *Rock Bass* | 24.9 | Cool | 2.56 | 3.04 | 2.05 | 1.02 | 1.72 |
| *Round Goby* | 20.7 | Cool | 0.26 | 0.10 | 0.34 | 0.34 | 0.69 |

| | | | | | | | |
|---|---|---|---|---|---|---|---|
| *Smallmouth Bass* | 25 | Warm | 16.12 | 26.91 | 32.14 | 17.69 | 9.97 |
| *Spoonhead Sculpin* | 6 | Cold | - | - | 0.17 | - | - |
| *Spottail Shiner* | 16.6 | Cold | 0.10 | - | 0.17 | 0.34 | - |
| *Threespine Stickleback* | 12.5 | Cold | 0.10 | - | - | - | - |
| *Trout-perch* | 13.4 | Cold | - | - | 0.17 | 0.34 | - |
| *Walleye* | 22.5 | Cool | 20.98 | 13.57 | 8.38 | 11.90 | 2.06 |
| *White Sucker* | 23.4 | Cool | 14.84 | - | 8.03 | 2.72 | 12.71 |
| *White Perch* | 29.8 | Warm | 1.54 | 0.94 | - | 0.68 | - |
| *White Bass* | 27.3 | Warm | - | 0.23 | - | - | - |
| *Yellow Perch* | 17.6 | Cool | 7.68 | 1.40 | 4.27 | 13.95 | 35.40 |

**Figure S1.** Map of sampling locations used in manuscript. Colour of site indicates what analysis data collected was used in. (SI = Stable Isotope Analysis; FA = Fatty Acid Analysis; Biomass = Biomass analysis using Broad-scale Monitoring Data). If sites were used for more than one analysis, all analysis used for are indicated by colour (e.g., data collected from Parry Sound was used for stable isotope, fatty acid and biomass analysis). Parry Sound is where the net-pen aquaculture facility is located and thus represents the experimental study site, all other sites are considered controls.

**Figure S2.** Comparison of the three % feed indicator fatty acids in net-pen feed to littoral forage fish (LFF), littoral baselines (LB), pelagic baselines (PB) and pelagic forage fish (PFF) between net-pen (Parry Sound) and control sites in 2016.

**Figure S3.** Comparison of the four % feed indicator fatty acids in net-pen feed to littoral forage fish (LFF), littoral baselines (LB), pelagic baselines (PB) and pelagic forage fish (PFF) between net-pen (Parry Sound) and control sites in 2017.

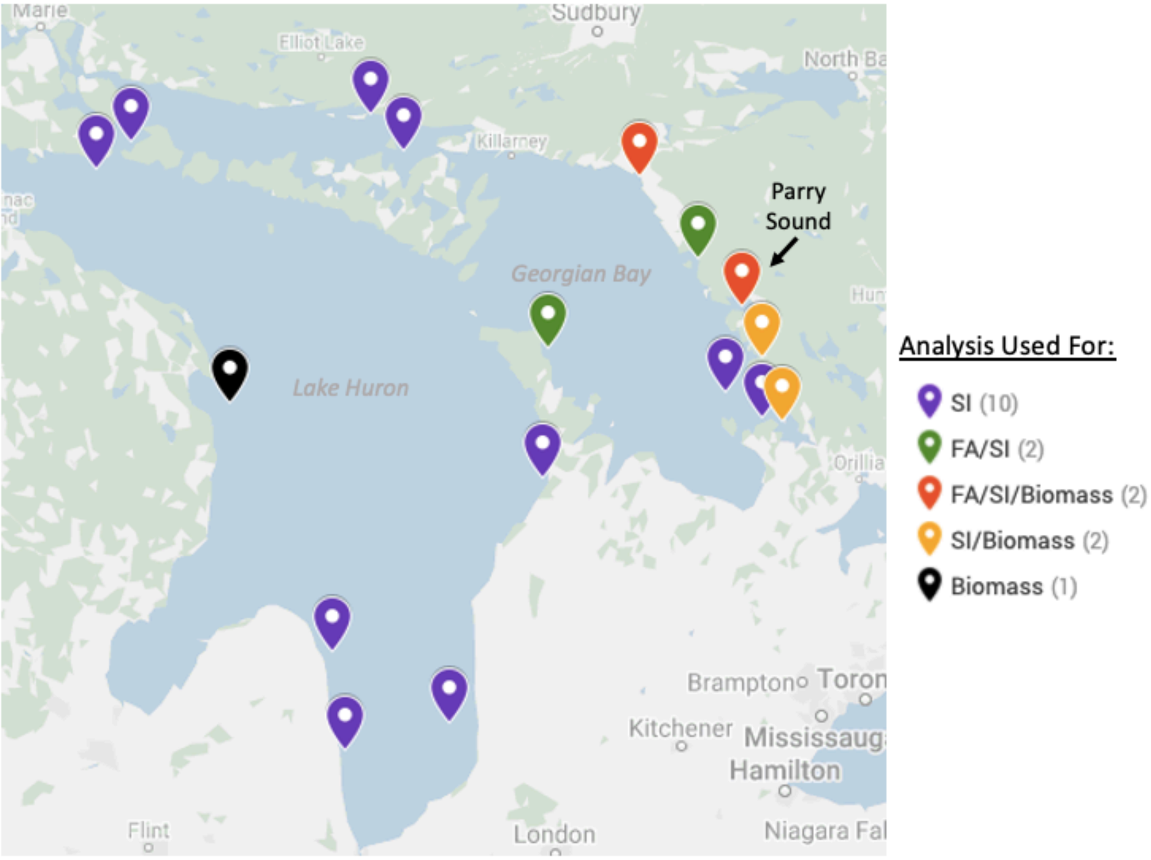
Marie
Sudbury
Elliot Lake
Killarney
Parry Sound
Georgian Bay
Lake Huron
Orillia
Brampton
Kitchener
Hamilton
Flint
London
Analysis Used For:
SI (10)
FA/SI (2)
FA/SI/Biomass (2)
SI/Biomass (2)
Biomass (1)

**Figure S1.**

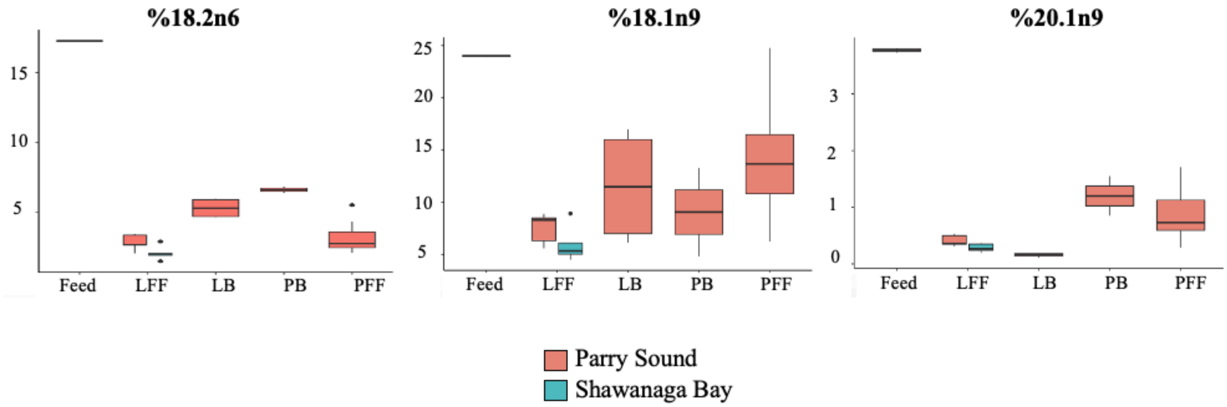

%18.2n6
%18.1n9
%20.1n9
Feed
LFF
LB
PB
PFF
Parry Sound
Shawanaga Bay


**Figure S2.**

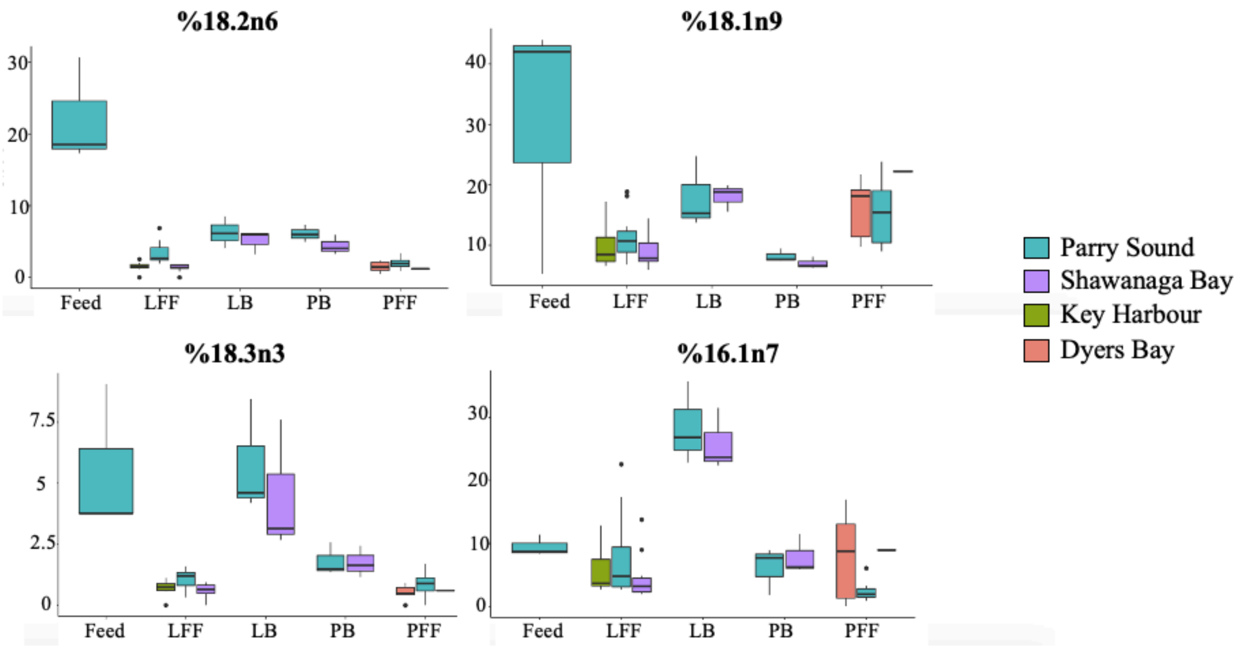
%18.2n6
30
20
10
0
Feed
LFF
LB
PB
PFF
%18.1n9
40
30
20
10
Feed
LFF
LB
PB
PFF
%18.3n3
7.5
5
2.5
0
Feed
LFF
LB
PB
PFF
%16.1n7
30
20
10
0
Feed
LFF
LB
PB
PFF
Parry Sound
Shawanaga Bay
Key Harbour
Dyers Bay

**Figure S3.**
